# Low-cost and open-source super-resolution fluorescence microscope with autofocus for teaching and research

**DOI:** 10.1101/2022.02.22.481481

**Authors:** Justin D. Hanselman, Benjamin G. Kopek

## Abstract

Super-resolution fluorescence microscopy revolutionized optical microscopy by breaking the theoretical resolution limit and thus providing an unprecedented view of biological structure in exquisite detail. Additionally, the relative ease of use and implementation for some super-resolution microscopes provided hope that this technology would be widely available and accessible. Despite initial excitement, super-resolution microscopy is still largely limited to facilities that can afford expensive commercial instruments or with personnel experienced in optical engineering that can construct an instrument. Recently, the open scientific hardware movement has attempted to make scientific instrumentation more widely available and accessible in a similar way to the open software movement. The widespread availability and affordability of prototyping devices, such as three-dimensional printers and interfaces (e.g., Arduino) have all contributed to unlocking the potential of the open hardware movement, turning expensive “black box” hardware into open-source, affordable, and robust instrumentation. Many others have recognized the potential of describing low-cost implementations of super-resolution microscope designs. However, we have found these microscope designs lacking in the documentation and details required to meet the requirements of open-source hardware. In this paper, we attempt to provide the documentation, details, and instructions necessary for the construction of an open-source and low-cost super-resolution microscope at a targeted cost of <$15,000.

## Introduction

Microscopes have been a main tool for uncovering cellular structure and function for centuries. The resolution of light microscopes was thought to be theoretically limited to approximately one-half the wavelength of emitted light (e.g., resolution limit of ∼250 nm for 500 nm wavelength light). However, multiple super-resolution microscopy methods have been developed that “break” the theoretical resolution limit and can achieve resolutions on the tens of nanometers scale in three dimensions[1–6]. These super-resolution fluorescence microscopes have revolutionized optical microscopy by providing an exquisitely detailed view of biological structure. Additionally, the relative ease of use and implementation for early super-resolution microscopes provided hope that this technology would be widely available, accessible, and relatively inexpensive. Despite this early promise, super-resolution microscopy is still largely limited to facilities that can afford expensive commercial instruments or with personnel experienced in optical engineering that can construct an instrument.

Recently, the open scientific hardware movement has sought to make scientific instrumentation more widely available and accessible in a similar way to the open software movement. The increasing availability and affordability of prototyping devices, such as three-dimensional printers and microcontrollers (e.g., Arduino) have helped the open hardware movement turn expensive “black box” hardware into open-source, affordable, and robust instrumentation[7–14]. This has begun to feed back into super-resolution microscopy and several researchers have described low-cost versions of super-resolution microscope designs[15–17]. However, we found these microscope designs lacking in the documentation and details required to meet open-source hardware standards[18]. In this paper, we attempt to provide the documentation, details, and instructions necessary for the construction of an open-source and low-cost super-resolution microscope.

The first consideration when building a super-resolution microscope is to determine which “method” is most suitable. New super-resolution methods are being rapidly developed and any list would soon be out of date. However, three methods for super-resolution microscopy that are currently the most popular and are commercially available include: (1) structured illumination microscopy (SIM), (2) STED (stimulated emission depletion) microscopy, and (3) single-molecule localization microscopy (SMLM). We will provide a brief overview of these various techniques and methods as it relates to our choice of the system developed, however, we refer the reader to the original articles and the many good reviews for a more in-depth discussion[1,19–35]. In particular, we recommend the review by Schermelleh et al. who provides a review that assists biologists in choosing the super-resolution method best-suited to their research question[35].

SIM is a combination of optical and image processing methods that can increase resolution approximately two-fold. This is accomplished by using striped patterns that produce Moiré fringes when interacting with features within the sample. Information is then decoded to reconstruct an image with increased resolution. Commercial systems exist that can be placed “on-top” of existing confocal systems, but there are several reasons why we did not choose a SIM microscope to construct. First, SIM relies on a sensitive (i.e., scientific grade camera), which increases the total system cost substantially. Secondly, the optical components for SIM require precise alignment and calibration, which if not performed properly, can result in reconstruction artifacts. Lastly, the resolution is only approximately two-fold. Still, SIM may be a good solution for many researchers, and we refer those interested to the following reference[36].

A second super-resolution technique is STED (stimulated emission depletion), where the shape of the PSF is engineered to smaller physical dimensions to produce higher resolution. This approach uses directed laser beams to achieve sub-diffraction limit resolutions (30-80 nm). STED requires simultaneous illumination of the sample with the confocal excitation beam overlaid by a depletion laser beam with a local intensity minimum in the focal center. This creates a “donut” shape where the size of the donut hole roughly corresponds to the resolution. Commercial add-ons are available that will allow one to simply perform STED on an existing confocal set-up. However, the optical components and lasers required to construct a STED system are complex and expensive, thus making a low-cost implementation more difficult.

Single-molecule localization microscopy (SMLM), includes PALM (photoactivated localization microscopy) and STORM (stochastic optical reconstruction microscopy). Sometimes referred to as “pointillist” approaches, these methods rely on the stochastic switching of individual fluorophores. The photons from a sparse subset of spatiotemporally separated fluorophores are collected and each molecule can then be localized with great precision. These methods are dependent on the properties of the fluorophores (i.e., fluorescent proteins, dyes), therefore, these methods require the least sophisticated microscope set-up of all super-resolution approaches and can be performed on a basic wide-field microscope. A drawback is that the sample must be prepared in a compatible way, such as using specific fluorescent proteins, dyes, and buffers. Overall, we have found that sample preparation can be simplified using standard dyes in a simple buffer system as first described by Olivier et al. and used for our results here[37]. If one is hardware limited then this approach becomes attractive on an ease-of-use and cost-effective basis. Thus, we describe here a microscope construction design that allows one to perform SMLM using a comparably simple design.

## Implementation

The overall design was based on the cost-efficient blueprint for localization microscopy created by Holm and colleagues[15], including an optical-cage system (Figure 1). Open optical-cage systems may be intimidating to those inexperienced in optical system construction, however, we found these to be no more difficult than putting together IKEA furniture and we wrote step-by-step assembly instructions with detailed component lists (available in the DocuBricks repository[38]). While custom parts were an inevitability, they were designed in such a way that standard 3-D printing or low-complexity machining is sufficient. Wherever possible, hardware was chosen to be a standard manufactured component from a large-scale retailer. Many of the solutions presented here are simply “what worked for us at the time” and not necessarily the optimal or perfect solution. In such cases, we will attempt to point out alternatives. As implemented, the estimated cost of all components is <$15,000, not including costs associated with personnel time.

**Figure 1.**
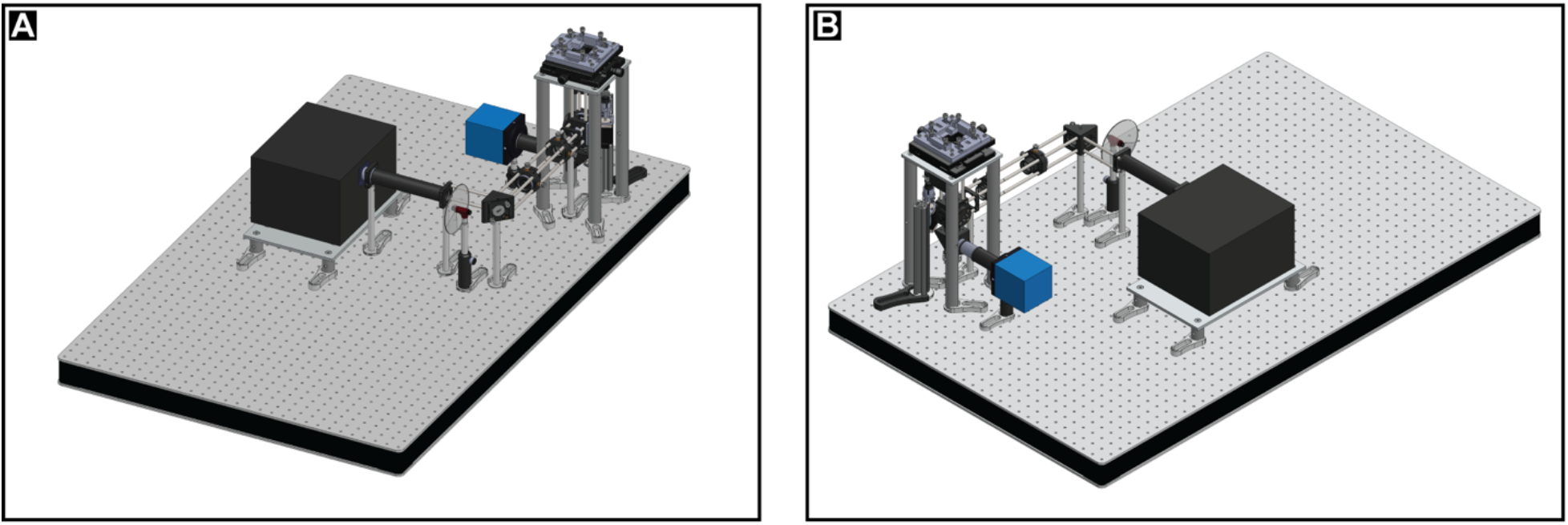
Overview of microscope. Details are provided in this paper and assembly instructions can be found in the hardware repository DocuBricks.

For construction of the optical-cage, we refer the reader to the step-by-step instructions provided in the accompanying Docubricks[38]. We attempted to purchase most of the materials from Thorlabs (www.thorlabs.com), which we generally found to have the lowest-cost and widest-selection of needed materials for the optical cage. All materials, including 3D printed and machined parts, are detailed and provided in Docubricks[38]. In the following sections, we discuss some of the major components (laser unit, camera, software, autofocus) and considerations and alternatives for each.

One major component that differentiates a super-resolution system from a more basic widefield microscope is the need for a high intensity light source (i.e., laser). Although some have reported using a Mercury Arc lamp for photoactivation and imaging[39], in general, a laser is going to be a better (i.e., faster) solution. Holm et al. made the innovative choice of using an entertainment grade laser[15]. Entertainment grade lasers can offer high-output at a lower cost compared to industrial or scientific grade lasers. Typically, different laser units would be used for each wavelength, but entertainment lasers have multiple lasers in a single unit. This may be an advantage or a disadvantage; an advantage because there are three laser lines for the price of one but a disadvantage because it is not easy to quickly switch between laser lines. The entertainment laser provides ∼300 mW for red (637 nm) and green (532 nm) laser lines and 700 mW for the blue (445 nm) laser line. The red wavelength of 637 nm is ideal for the dye Alexa Fluor 647, which is considered the best dye available for direct Stochastic Optical Reconstruction Microscopy (dSTORM)[40]. The red laser is the ideal choice for single color applications and is the only laser we imaged with here. The green laser wavelength is 532 nm, which is not optimal for the excitation of many commonly used dyes and photo-activatable fluorescent proteins (PA-FP). However, the popular PA-FP mEos has an excitation shoulder at 532 nm and photoconverts at 532 nm as well, making the localization of mEos tagged proteins possible with this laser system as demonstrated by Holm et al.[15]. Another fluorescent dye that could be used in this setup is cy3B, which also has a shoulder at ∼532 nm, and thus could be imaged with our setup[41]. Also, several blue dyes (Atto 488) and fluorescent proteins (PA-GFP) could be excited using the 455 nm laser, though this remains untested in our system. Photoactivation of PA-FPs is typically done using simultaneous illumination with a 405 nm laser. This microscope does not currently allow for simultaneous laser illumination; however, a separate 405 nm laser could be added. One can purchase lower-cost laser units, especially previously used and industrial-grade units, as an alternative to the entertainment laser unit used here.

An area that is not typically discussed in publications on low-cost microscope construction is laser control. Scientific and industrial grade lasers may come with software supplied by the manufacturer. However, because we opted for a lower-cost entertainment-grade laser, we had to construct a circuit board to interface with the International Laser Display Association (ILDA) laser control standard. Although we initially used a breadboard for prototyping, a circuit board could be printed at minimal cost (schematics are available in Docubricks file[38]). We then wrote a laser control plugin for the microscope management software (plugin available on GitHub[42]). The result was a high-powered, lower-cost illumination source that was easily controllable.

Another critical component of an SMLM microscope is the camera. The ability to use a camera versus a detector for SMLM makes microscope construction easier compared to the other super-resolution based approaches that rely on point-scanning (e.g, STED). A high quantum efficiency (number of photoelectrons generated per incident photon) is required to maximize the number of photons detected since localization precision is dependent on the inverse square root of the number of collected photons (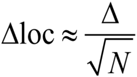; where Δloc is the localization precision, Δ is the FWHM of the PSF and *N* is the number of collected photons)[43]. Therefore, obtaining the highest resolution requires an electron multiplier charged-coupled device (EMCCD) camera. High-performance EMCCD cameras have a quantum efficiency >90%, while standard CCD cameras typically have a quantum efficiency around 40-50%. However, the quantum efficiency of inexpensive CCD cameras continues to improve, and many have a quantum efficiency >75% in portions of the visible spectrum. For our system, we chose the Retiga R3 from Q imaging. The Retiga R3 features a quantum efficiency of 75% at 600 nm and a pixel size of 4.54 um x 4.54 um at a cost of $5,463. The cost of the camera accounts for nearly a third the cost of the entire system, however, this is still a sizable savings compared to an EMCCD camera with a cost of >$30,000. Another solution for the reader to consider is the CMOS camera used by Ma et al.[16]. The CMOS camera sensor is a lower cost (<$500) solution that could be adapted to our setup and should be considered for any low-cost system. We had already purchased our camera when we became aware of the CMOS camera solution and our camera came with acquisition software. As camera technology continues to improve with costs decreasing relative to quality, many good options will arise for implementation in SMLM.

Algorithmically reconstructing an SMLM image requires acquisition of tens of thousands of camera frames resulting in lengthy acquisition times (30 minutes to 1 hour). In our initial testing, we found that the microscope was only capable of staying in focus for ∼5-10 minutes. Maintaining a sample in focus for longer periods of time (tens of minutes to an hour) is often difficult even on systems that are mechanically stable. Mechanical and thermal fluctuations and instability all contribute to focus drifts and even the opening of a room door or air movement within the room can result in substantial drift. While one could continuously observe the image and adjust focus manually, this quickly becomes untenable. Therefore, it became necessary to find a solution for keeping the region of interest in focus over the time scales required for SMLM acquisition.

Many strategies for autofocusing have been developed but can be broadly categorized as software focusing algorithms such as[44–46] or optical approaches such as[47,48]. For both categories, movement of the stage or objective must occur to correct for drift during image acquisition. Commonly, such movement is done using a piezoelectric stage or objective actuator. Movement generated by piezoelectric materials is often ideal as it allows for very small movements. However, when considering our options, piezoelectric hardware was beyond our limited funds. Therefore, we designed an autofocus system using a low-cost motor (∼$500) that drives objective movement. Focus is detected using markers (e.g., gold nanoparticles). Initial focus is set by the user and software was written to refocus through movement of the objective using the markers to gauge whether focus is achieved.

The open-source microscope control software Micromanager (μManager) was used to provide control of the laser, camera, and autofocus motor[49]. Camera control was straightforward as the Retiga R3 is μManager compatible and many controls are included in the μManager software. For laser and autofocus control using μManager, several plugins were written (available on GitHub[42]). A Microsoft Windows-based desktop computer running Windows 7 with a 3.4 GHz Intel Xeon processor, 32 GB RAM, and 1 TB hard drive was used to run the hardware, for image acquisition, and data processing.

The output from the acquisition is a series of Tiff files. Software was written to initially correct for drift. To process the images (localize single molecules), a variety of software solutions are available and reviewed at http://bigwww.epfl.ch/smlm/software/[50,51]. Our currently preferred software solution is Thunderstorm, which exists as an ImageJ plugin[52].

## Safety

The illumination source is a high-powered entertainment laser and all appropriate laser safety precautions should be taken, including the use of laser safety glasses and beam blocks. We used a beam block that extends out from the laser aperture and a laser safety box should be used during setup and operation to block errant beams. Alignment of the laser poses the greatest risk of laser exposure and appropriate laser safety glasses are recommended. To align the laser, we found that an inexpensive camera (e.g., webcam) can be installed to help guide alignment. During illumination of the sample, the high-powered laser beam exits the objective and projects through the sample and onto the ceiling. An enclosure can be built above and around the stage to prevent errant beams from reflective surfaces. We recommend the purchase of commercially available optical enclosures.

While the laser is the most obvious source of danger, working with electronics and electronic components poses the risk of electrical shock. Standard electrical safety procedures should be followed if the power-supply box is constructed.

### Calibration

There are many software solutions available for SMLM processing data and providing a measure of resolution or localization precision (http://bigwww.epfl.ch/smlm/software/)[50]. We prefer the straightforward measurement of “Full Width at Half Maximum (FWHM).” Resolution in SMLM is dependent on the density of fluorescent labels and the structure of the sample, thus we recommend testing the microscope on a sample of known size (e.g., beads, cytoskeletal elements). Others have documented ways to determine resolution and image quality in SMLM including[53] and[54].

### General testing

Super-resolution fluorescence microscopes are primarily intended for the localization of fluorescent dyes or proteins in biological samples. However, other samples can be imaged provided that the spectral properties meet the requirements (i.e., appropriate wavelength, ability to photo-activate or photo-switch). For this set-up, we recommend Alexa Fluor 647 labeled samples imaged using dSTORM. dSTORM requires special buffers, however, Vectashield mounting medium is a simple and effective solution[37].

A sample that can be used for general microscope testing is pre-mounted fluorescent beads (e.g. TetraSpeck; ThermoFisher #T7279) that come in a variety of sizes and excitation wavelengths allowing the user to test alignment, focus, and overall microscope function. For testing of the autofocus system proposed here, the use of gold fiducial nanoparticles is recommended. The nanoparticles can be mounted on a glass coverslip or chambered coverslip. It is important that the appropriate size of nanoparticle is selected for the wavelength tested.

Testing the super-resolution capabilities of the microscope can be done using generic cells that are labeled with commercially available dyes. We recommend using phalloidin conjugated to Alexa Fluor 647 (see Application), which binds strongly to actin filaments. Many protocols for actin labeling with phalloidin are available, including the one described here. Because of the thin filamentous nature of cytoskeletal components, they are ideal test cases to gauge system resolution.

### Application

#### Use case(s)

For testing purposes, we imaged the cytoskeletal elements actin and tubulin in Vero cells (*Cercopithecus aethiops* kidney epithelial cells; ATCC CCL-81). Vero cells are a convenient cell culture system because they are flat and adherent.

Fiducial markers must be deposited on the glass surface for drift correction and autofocus. We used 8-well chambered coverslips (Lab-Tek II Chambered Coverglass #1.5 borosilicate, Thermo Fisher Sci. #155409) because cells can be plated and grown in them and the chamber allows for the addition of a buffer during imaging. To add gold fiducials to the coverslip, 200 ul of a 0.01% poly-L-lysine solution (Millipore-Sigma #P4832) was added to each well and allowed to incubate at room temperature for 10 minutes. After the incubation, the poly-L-lysine solution was aspirated from the wells. Gold Nanorods 25 nm in diameter and 71 nm in length (Nanopartz #A12-25-650-25-CTAB-DI) appropriate for 637 nm excitation were diluted in sterile deionized water. An appropriate dilution (one that provides enough particles for drift correction and autofocus; typically, 5-8 particles within a field-of-view) should be determined empirically as the gold particles are a colloid that can settle and clump. In general, a dilution of 1:100 to 1:200 is appropriate but sometimes a 1:1 dilution was needed to get sufficient markers. We are unable to determine if a “spot” represents a single nanorod or multiple nanorods, regardless, these function as a single point. After poly-L-lysine coating, we added 200 ul of diluted nanorods to the coverslip and incubated for 15 minutes at room temperature. Nanorods were aspirated and the coverslips were washed once with sterile deionized water before aspiration of all liquid.

Vero cells were added to appropriate wells of the chambered coverglass at 1 × 10^4^ cells per well in 400 µL of medium. Actin was labeled with an Alexa Fluor 647-phalloidin conjugate (ThermoFisher #A22287) as previously described[55,56], although a detailed protocol is provided here as well. After an overnight incubation at 37 °C in 5% CO_2_ to allow the cells to attach and grow, the cells were fixed and permeabilized for 2 minutes with 0.3% glutaraldehyde with 0.25% triton X-100 in Cytoskeleton Buffer (CB: 10 mM MES pH 6.1, 150 mM NaCl, 5 mM EDTA, 5 mM glucose, and 5 mM MgCl_2_). A second fixation was performed using 2% glutaraldehyde in CB for 10 minutes. To reduce autofluorescence caused by aldehydes, the samples were treated with 0.1% sodium borohydride in 100 mM phosphate buffer for seven minutes followed by three, 10-minute washes, with phosphate buffered saline pH 7.4 (PBS). Samples were blocked with 5% bovine serum albumin (BSA) in PBS for 30 minutes. The Alexa Flour 647-phalloidin conjugate was diluted 1:40 in 5% BSA-PBS and incubated with the cells overnight at 4 °C. After the overnight incubation with the dye, the cells were washed with 0.1% Tween-20 in PBS for 2 minutes followed by a PBS wash for 2 min. The PBS was aspirated and soft-mount Vectashield (H-1000) was added to the cells for storage and imaging.

An example of the images of labeled actin using this system can be seen in Figure 2. This image is qualitatively on par with our previous images of actin using more advanced systems[55,56].

**Figure 2.**
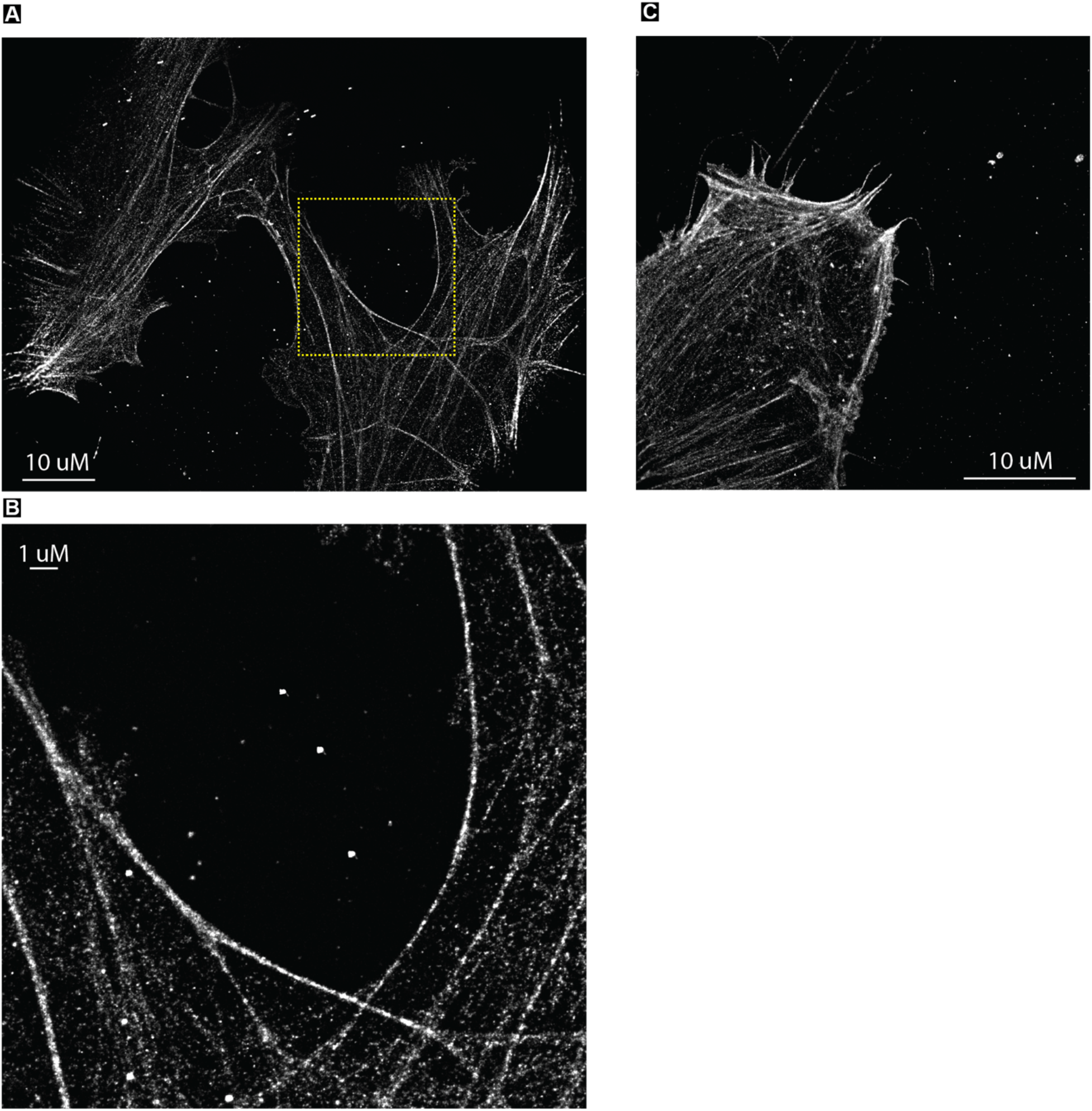
Super-resolution image of actin filaments. (A) Vero cells were labeled with phalloidin conjugated to Alexa Fluor 647. (B) Magnified view of box in (A). There are multiple fiducials (dots) in the middle of the image. (C) Another example of a labeled cell.

Antibody labeling allows a large number of targets to be imaged with this system. Here the cytoskeletal element tubulin was labeled using an anti-tubulin primary antibody (Abcam #ab18251) and a secondary antibody conjugated to Alexa Fluor 647 (ThermoFisher #A-21244). A similar method of sample preparation was used for tubulin labeling as was done for actin labeling. As with actin labeling, the samples were fixed and permeabilized, treated with sodium borohydride, and washed with PBS. Blocking was done with a solution of 5% BSA and 10% normal goat serum (NGS) in PBS for 30 minutes at room temperature. The anti-tubulin antibody was diluted to a concentration of 1 ug/ml in a 5% BSA in PBS solution. Cells were incubated with the anti-tubulin antibody overnight at 4 °C. After the primary antibody incubation, cells were washed three times with PBS for 5 min each. Alexa Fluor 647 labeled goat anti-rabbit secondary antibody was diluted to a concentration of 4 ug/mL in 5% BSA in PBS solution and incubated with the cells for 1 hour at room temperature protected from light. Following the secondary antibody incubation, cells were washed three times with PBS for 5 minutes each, one time with 0.1% Tween-20 in PBS for 2 minutes, and once with PBS for 2 minutes. Vectashield was added to the chambers for storage and imaging. An example of cells labeled for tubulin in this manner can be seen in Figure 3.

**Figure 3.**
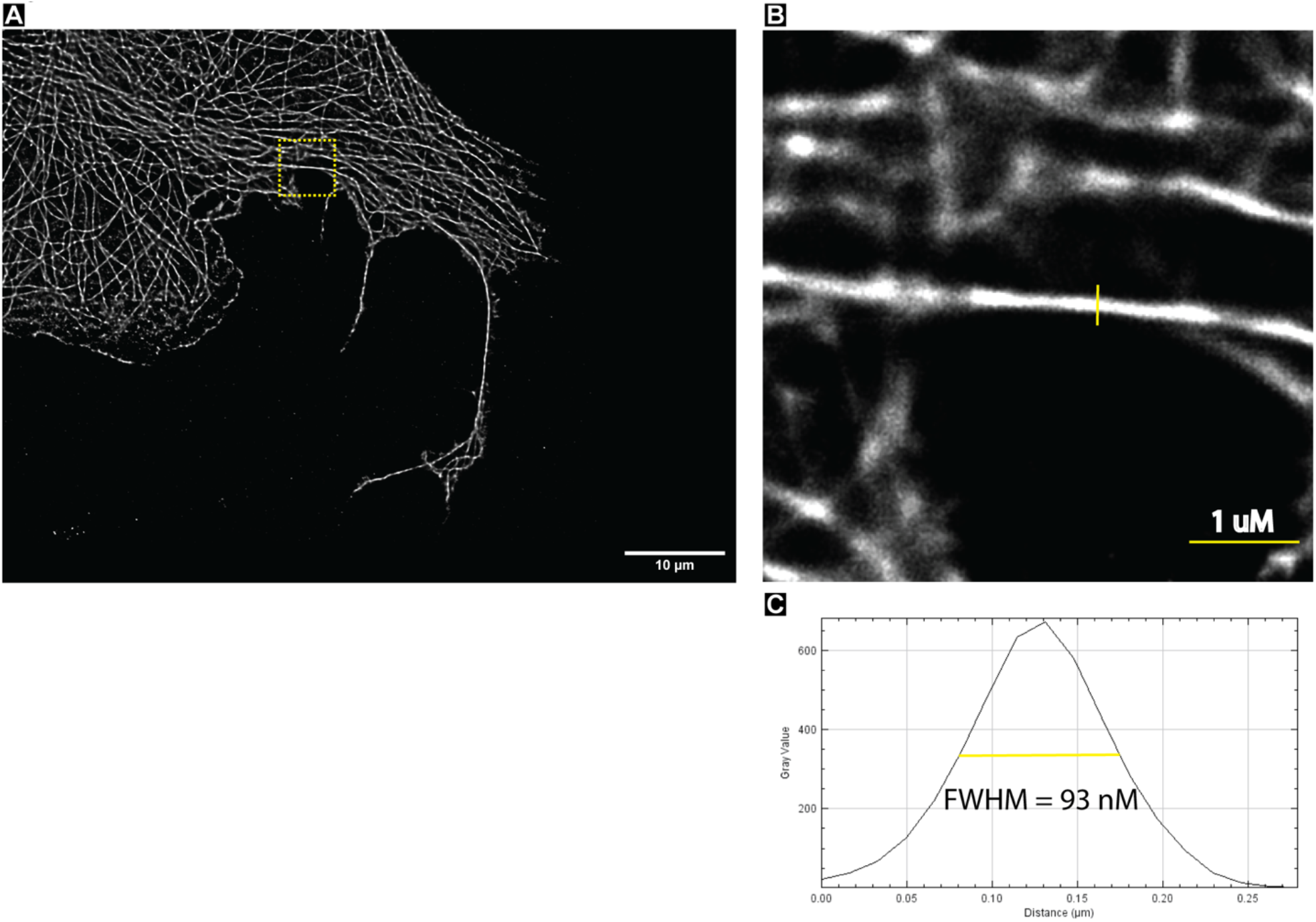
Super-resolution image of microtubules. (A) Vero cells were labeled with anti-tubulin antibodies and secondary antibodies conjugated to Alexa Fluor 647. (B) Magnified view of box in (A). (C) Line plot showing the FWHM = 93 nm of the line through the microtubule in (B). Many super-resolution techniques show a FWHM of ∼50-60 nm for microtubules[37].

The autofocus system was robust, allowing acquisitions over 30,000 frames (over an hour of acquisition time). This was more than sufficient to achieve well-resolved images. Longer acquisition times are possible, but we are limited by the memory of the computer (30 k frames is 20 GB of data that are stored in the system’s RAM). Sub-diffraction localization was done in Thunderstorm[52] using sub-pixel localization using radial symmetry[57], which was faster and sufficient for our processing needs.

#### Reuse potential and adaptability

Super-resolution images of Alexa Fluor 647 labeled samples using dSTORM were obtained. However, this system would allow for the imaging of other dyes, and fluorescent proteins, such as Atto 488 or mEos derivatives, using various SMLM techniques such as PALM and STORM. Additional laser lines can be added to allow for a greater variety of fluorescent targets, though this would require modification of the beam path and additional control software. Micromanager software provides the ability to control additional lasers.

Although the autofocus and drift correction presented here is a robust and low-cost solution, other autofocus systems could easily be implemented, such as a Perfect Focus System (Nikon). A fiducial-based z-focusing system using a piezio electric stage was implemented by Ma et al.[58].

These SMLM techniques are based on wide-field microscopes, thus limiting z-resolution. However, better z-resolution could be obtained using a total internal reflection (TIRF) objective, but TIRF imaging is limited to ∼< 200 nm above the coverglass, requiring thin samples or targets near the glass surface. Highly Inclined and Laminated Optical sheet (HiLO) is another method for improving z-resolution in super-resolution microscopy[59]. 3D SMLM modalities that could be implemented at a low cost include: astigmatic imaging[2], biplane imaging[60], and double helix point spread function microscopy[61].

A wide-variety of sophisticated 3D imaging approaches including dual-objective solutions like iPALM[1], beam sculpting[27], and Tilt3D[25] could also be considered, however at an increased cost.

### Build Details

#### Availability of materials and methods

The majority of optical cage components are commercially available through Thorlabs. We purchased the stage through Owis but equivalents are available on Thorlabs. When purchasing the stage, one should consider the types of samples that will be imaged. The camera used here was purchased through QImaging, though many other options for cameras exist. The laser unit was purchased online through Laserworld. The Olympus objective was purchased through a local Olympus dealer, but an equivalent objective is available from every major microscope manufacturer (Nikon, Zeiss). Optical filters were purchased from Edmund Optics. A variety of other parts were purchased through various commercial vendors, including Oriental Motors. Many of the other parts can be purchased locally. Although we constructed a laser safety box using components from a local hardware store, these are also available through Thorlabs, which we recommend for ease of construction. Several 3D printed parts and machined parts were used. Detailed assembly instructions with materials are included in the associated DocuBricks[38].

#### Ease of build

We have provided the build documentation and software to make construction straightforward. More difficult aspects include the wiring of a breadboard control for the laser unit. However, the target audience of undergraduate educators or biologists should be able to construct this system relatively easily.

#### Operating software and peripherals

The main software used here is Micromanager. Micromanager integrates with the camera and laser through a custom electrical controller. A device plugin was written in C++ to interface the electrical controller with micromanager (https://github.com/HopeCollegeSuperresolutionMicroscope). Additionally, a java plugin was written to enable fiducial tracking and autofocus behavior (https://github.com/HopeCollegeSuperresolutionMicroscope). An additional device plugin was written in C++ to interface a stepper motor and its controller with micromanager in order to automate focusing.

Micromanager software is available for both Windows and MacOS computers. We used a computer running Windows 7. Data sets of 10-20 GB are produced during image acquisition and stored in RAM. Therefore, we recommend a minimum of 32 GB RAM for most applications. A large (1 TB or more) hard drive for data storage is an obvious benefit as is a multi-core processor. These specifications are a minimum for most applications and better computer hardware specifications are desirable.

### Hardware documentation and files location

***Archive for hardware documentation and build files***

***Name:*** DocuBricks

***Persistent identifier:***http://docubricks.com/viewer.jsp?id=7979244493125740544

***Licence:*** CC BY 3.0

***Publisher:*** B.G. Kopek

***Date published:*** 11/09/19

**Software code repository**

***Name:*** GitHub

***Identifier:*** https://github.com/HopeCollegeSuperresolutionMicroscope

***Licence:*** CC BY 3.0

***Date published:*** 12/10/20

## Discussion

The initial introduction of super-resolution fluorescence microscopy was hailed for its relative low-cost and simplicity. For “less than the price of a new kitchen” someone could build a super-resolution microscope[62]. In fact, one of the first super-resolution microscopes was built in an apartment with off-the-shelf parts. However, these technologies were quickly licensed to companies for commercialization. While commercial systems can provide a user-friendly interface there is a trade-off in an increased cost, which leaves such systems not available to many researchers. Commercial fluorescence microscope systems typically cost in the hundreds of thousands of US dollars while a self-constructed system can be in the thousands to a few tens of thousands of US dollars. The microscope described here has a cost of ∼$15,000 while producing images with sub-diffraction limit resolution. Overall, this system should provide a good starting reference for the construction of a low-cost super-resolution fluorescence microscope system. We believe our low-cost auto-focus system to be a major contribution to the field as other solutions add significant cost to the system. However other, likely easier, implementations of drift correction and autofocus could be used if cost allows. Additional laser lines could be added with minor modification, yet typical super-resolution experiments are a single color and make use of Alexa Fluor 647. Thus, this system will produce data for these experiments without the cost and complexity of additional lasers.

In our opinion, an often-overlooked area of need for low-cost microscopy is undergraduate education. Liberal arts colleges (which are typically primarily undergraduate institutions) in the USA train a disproportionate number of students who go on to earn doctorate degrees in science and engineering compared to research universities[63]. However, primarily undergraduate institutions often lack the user base large enough to warrant acquisition of a commercial super-resolution system or the funds and expertise to construct a home-built system. Yet science is becoming more interdisciplinary, requiring researchers to work across traditional disciplines. Microscopy requires knowledge in engineering, physics, chemistry (e.g., dyes), biology, and computer science. If educators at primarily undergraduate institutions felt enabled to construct simple, yet powerful microscopes at a reasonable cost it may open new avenues in interdisciplinary education. Lastly, many researchers in the developing world lack resources for advanced microscopy putting themselves and their countries at a severe disadvantage. Providing ways for these researchers to increase their research productivity at a low cost will aid the democratization of science.

As stated previously, the system described here is not meant to be an endpoint, finalized solution. However, we hope the information provided will make construction and use of super-resolution fluorescence microscopes easier for anyone who wishes to do so. Open hardware solutions for scientific instrumentation have the potential to save the scientific community large amounts of money and aid the democratization of science.

## Paper author contributions

Design, assembly, documentation, paper writing – J.H.

Conceived idea, use cases contribution, documentation, paper writing – B.G.K.

## Acknowledgements

We thank other Kopek Lab members that assisted with sample preparation including Haley Fischman.

## References

1. Shtengel G, Galbraith JA, Galbraith CG, Lippincott-Schwartz J, Gillette JM, Manley S, et al. Interferometric fluorescent super-resolution microscopy resolves 3D cellular ultrastructure. Proc National Acad Sci. 2009;106: 3125–3130. doi:10.1073/pnas.0813131106

2. Huang B, Jones SA, Brandenburg B, Zhuang X. Whole-cell 3D STORM reveals interactions between cellular structures with nanometer-scale resolution. Nat Methods. 2008;5: 1047–1052. doi:10.1038/nmeth.1274

3. Betzig E, Patterson GH, Sougrat R, Lindwasser OW, Olenych S, Bonifacino JS, et al. Imaging intracellular fluorescent proteins at nanometer resolution. Science. 2006;313: 1642–1645. doi:10.1126/science.1127344

4. Hofmann M, Eggeling C, Jakobs S, Hell SW. Breaking the diffraction barrier in fluorescence microscopy at low light intensities by using reversibly photoswitchable proteins. Proc Natl Acad Sci U S A. 2005;102: 17565–17569. doi:10.1073/pnas.0506010102

5. Hess ST, Girirajan TPK, Mason MD. Ultra-high resolution imaging by fluorescence photoactivation localization microscopy. Biophysical Journal. 2006;91: 4258–4272. doi:10.1529/biophysj.106.091116

6. Rust MJ, Bates M, Zhuang X. Sub-diffraction-limit imaging by stochastic optical reconstruction microscopy (STORM). Nat Methods. 2006;3: 793–795. doi:10.1038/nmeth929

7. Banović L, Vihar B. Development of an Extruder for Open Source 3D Bioprinting. J Open Hardw. 2018;2: 1. doi:10.5334/joh.6

8. Delmans M, Haseloff J. μCube: A Framework for 3D Printable Optomechanics. J Open Hardw. 2018;2: 2. doi:10.5334/joh.8

9. Wayland MT, Landgraf M. A Cartesian Coordinate Robot for Dispensing Fruit Fly Food. J Open Hardw. 2018;2: 3. doi:10.5334/joh.9

10. McGinness L, Dührkoop S, Jansky A, Keller O, Lorenz A, Schmeling S, et al. 3D-Printable Model of a Particle Trap: Development and Use in the Physics Classroom. J Open Hardw. 2019;3: 1. doi:10.5334/joh.12

11. Muschard B, Bonvoisin J. CubeFactory2 – an Off-Grid and Circular 3D-Printing Mini-Factory. J Open Hardw. 2019;3: 3. doi:10.5334/joh.15

12. Conley K, Foyer A, Hara P, Janik T, Reichard J, D’Souza J, et al. Vibration Alert Bracelet for Notification of the Visually and Hearing Impaired. J Open Hardw. 2019;3: 4. doi:10.5334/joh.17

13. Bowman RW, Vodenicharski B, Collins JT, Stirling J. Flat-Field and Colour Correction for the Raspberry Pi Camera Module. J Open Hardw. 2020;4: 1. doi:10.5334/joh.20

14. Jonveaux L. Arduino-like development kit for single-element ultrasound imaging. J Open Hardw. 2017;1: 3. doi:10.5334/joh.2

15. Holm T, Klein T, Klamp T, Löschberger A, Wiebusch G, Wiebusch G, et al. A Blueprint for Cost-Efficient Localization Microscopy. ChemPhysChem. 2013;15. doi:10.1002/cphc.201300739

16. Ma H, Fu R, Xu J, Liu Y. A simple and cost-effective setup for super-resolution localization microscopy. Scientific Reports. 2017;7: 1–9. doi:10.1038/s41598-017-01606-6

17. Kwakwa K, Savell A, Davies T, Munro I, Parrinello S, Purbhoo MA, et al. easySTORM: a robust, lower-cost approach to localisation and TIRF microscopy. Journal of biophotonics. 2016;9: 948–957. doi:10.1002/jbio.201500324

18. Definition (English) – Open Source Hardware Association. Available: https://perma.cc/F4UN-787X

19. Bates M, Huang B, Dempsey GT, Zhuang X. Multicolor super-resolution imaging with photo-switchable fluorescent probes. Science. 2007;317: 1749–1753. doi:10.1126/science.1146598

20. Dertinger T, Colyer R, Iyer G, Weiss S, Enderlein J. Fast, background-free, 3D super-resolution optical fluctuation imaging (SOFI). Proc Natl Acad Sci U S A. 2009;106: 22287– 22292. doi:10.1073/pnas.0907866106

21. Dertinger T, Pallaoro A, Braun G, Ly S, Laurence TA, Weiss S. Advances in superresolution optical fluctuation imaging (SOFI). Q Rev Biophys. 2013;46: 210–221. doi:10.1017/s0033583513000036

22. Carlini L, Manley S. Live Intracellular Super-Resolution Imaging Using Site-Specific Stains. ACS Chemical Biology. 2013;8: 2643–2648. doi:10.1021/cb400467x

23. Fornasiero EF, Opazo F. Super-resolution imaging for cell biologists: Concepts, applications, current challenges and developments. BioEssays. 2015; n/a-n/a. doi:10.1002/bies.201400170

24. Gustafsson N, Culley S, Ashdown G, Owen DM, Pereira PM, Henriques R. Fast live-cell conventional fluorophore nanoscopy with ImageJ through super-resolution radial fluctuations. Nat Commun. 2016;7: 12471. doi:10.1038/ncomms12471

25. Gustavsson A-K, Petrov PN, Lee MY, Shechtman Y, Moerner WE. 3D single-molecule super-resolution microscopy with a tilted light sheet. Nature communications. 2018;9: 123. doi:10.1038/s41467-017-02563-4

26. Heilemann M, Linde S van de, Mukherjee A, Mukherjee A, Sauer M. Super-Resolution Imaging with Small Organic Fluorophores. Angewandte Chemie International Edition. 2009;48: 6903–6908. doi:10.1002/anie.200902073

27. Jia S, Vaughan JC, Zhuang X. Isotropic three-dimensional super-resolution imaging with a self-bending point spread function. Nature photonics. 2014;8: 302–306. doi:10.1038/nphoton.2014.13

28. Kaufmann R, Schellenberger P, Seiradake E, Dobbie IM, Jones EY, Davis I, et al. Super-resolution microscopy using standard fluorescent proteins in intact cells under cryo-conditions. Nano Letters. 2014;14: 4171–4175. doi:10.1021/nl501870p

29. Lippincott-Schwartz J, Manley S. Putting super-resolution fluorescence microscopy to work. Nat Methods. 2009;6: 21–23. doi:10.1038/nmeth.f.233

30. Manley S, Gunzenhäuser J, Olivier N. A starter kit for point-localization super-resolution imaging. Current Opinion in Chemical Biology. 2011;15: 813–821. doi:10.1016/j.cbpa.2011.10.009

31. Nanguneri S, Flottmann B, Horstmann H, Heilemann M, Kuner T. Three-dimensional, tomographic super-resolution fluorescence imaging of serially sectioned thick samples. PloS One. 2012;7: e38098. doi:10.1371/journal.pone.0038098

32. Schermelleh L, Heintzmann R, Leonhardt H. A guide to super-resolution fluorescence microscopy. J Cell Biol. 2010;190: 165–175. doi:10.1083/jcb.201002018

33. Wicker K. Super-Resolution Fluorescence Microscopy Using Structured Illumination. Humana Press; 2014. pp. 133–165. doi:10.1007/978-1-62703-983-3_7

34. Oijen AMV, Köhler J, Schmidt J, Müller M, Brakenhoff GJ. 3-Dimensional super-resolution by spectrally selective imaging. Chemical Physics Letters. 1998;292: 183–187. doi:10.1016/s0009-2614(98)00673-3

35. Schermelleh L, Ferrand A, Huser T, Eggeling C, Sauer M, Biehlmaier O, et al. Super-resolution microscopy demystified. Nat Cell Biol. 2019;21: 72–84. doi:10.1038/s41556-018-0251-8

36. Murphy DB, Davidson MW. Fundamentals of Light Microscopy. Fundamentals of Light Microscopy and Electronic Imaging. John Wiley & Sons, Ltd; 2012. pp. 1–19. doi:10.1002/9781118382905.ch1

37. Olivier N, Keller D, Rajan VS, Gönczy P, Manley S. Simple buffers for 3D STORM microscopy. Biomedical Optics Express. 2013;4: 885–15. doi:10.1364/boe.4.000885

38. Kopek BG, Hanselman JD. DocuBricks - Super-resolution Fluorescence Microscope. [cited 1 Jun 2020]. Available: https://www.docubricks.com/viewer.jsp?id=7979244493125740544

39. Prakash K. High-density superresolution microscopy with an incoherent light source and a conventional epifluorescence microscope setup. Biorxiv. 2019; 121061. doi:10.1101/121061

40. Linde S van de, Klein T, Löschberger A, Heidbreder M, Wolter S, Heilemann M, et al. Direct stochastic optical reconstruction microscopy with standard fluorescent probes. Nat Protoc. 2011;6: 991–1009. doi:10.1038/nprot.2011.336

41. Cooper M, Ebner A, Briggs M, Burrows M, Gardner N, Richardson R, et al. Cy3B™: Improving the Performance of Cyanine Dyes. J Fluoresc. 2004;14: 145–150. doi:10.1023/b:jofl.0000016286.62641.59

42. Hanselman JD. HopeCollegeSuperresolutionMicroscope † GitHub. [cited 22 Feb 2022]. Available: https://github.com/HopeCollegeSuperresolutionMicroscope

43. Mortensen KI, Churchman LS, Spudich JA, Flyvbjerg H. Optimized localization analysis for single-molecule tracking and super-resolution microscopy. Nat Methods. 2010;7: 377–381. doi:10.1038/nmeth.1447

44. Price JH, Gough DA. Comparison of phase-contrast and fluorescence digital autofocus for scanning microscopy. Cytometry. 1994;16: 283–297. doi:10.1002/cyto.990160402

45. Li S, Cui X, Huang W. High resolution autofocus for spatial temporal biomedical research. Rev Sci Instrum. 2013;84: 114302. doi:10.1063/1.4829616

46. Yazdanfar S, Kenny KB, Tasimi K, Corwin AD, Dixon EL, Filkins RJ. Simple and robust image-based autofocusing for digital microscopy. Opt Express. 2008;16: 8670. doi:10.1364/oe.16.008670

47. Liron Y, Paran Y, Zatorsky NG, Geiger B, Kam Z. Laser autofocusing system for high-resolution cell biological imaging. J Microsc-oxford. 2006;221: 145–151. doi:10.1111/j.1365-2818.2006.01550.x

48. Pertsinidis A, Zhang Y, Chu S. Subnanometre single-molecule localization, registration and distance measurements. Nature. 2010;466: 647–651. doi:10.1038/nature09163

49. Stuurman N, Edelstein A, Amodaj N, Hoover K, Vale R. Computer control of microscopes using µManager. Curr Protoc Mol Biology. 2010;Chapter 14:Unit14.20. doi:10.1002/0471142727.mb1420s92

50. Sage D, Kirshner H, Pengo T, Stuurman N, Min J, Manley S, et al. Quantitative evaluation of software packages for single-molecule localization microscopy. Nature Methods. 2015;12: 717– 724. doi:10.1038/nmeth.3442

51. Sage D, Pham T-A, Babcock H, Lukes T, Pengo T, Chao J, et al. Super-resolution fight club: Assessment of 2D & 3D single-molecule localization microscopy software. Nat Methods. 2019;16: 387–395. doi:10.1038/s41592-019-0364-4

52. Ovesný M, Křížek P, Borkovec J, Švindrych Z, Hagen GM. ThunderSTORM: a comprehensive ImageJ plug-in for PALM and STORM data analysis and super-resolution imaging. Bioinformatics. 2014;30: 2389–2390. doi:10.1093/bioinformatics/btu202

53. Nieuwenhuizen RPJ, Lidke KA, Bates M, Puig DL, Grünwald D, Stallinga S, et al. Measuring image resolution in optical nanoscopy. Nature Methods. 2013;10: 557–562. doi:10.1038/nmeth.2448

54. Culley S, Albrecht D, Jacobs C, Pereira PM, Leterrier C, Mercer J, et al. Quantitative mapping and minimization of super-resolution optical imaging artifacts. Nat Methods. 2018;15: 263–266. doi:10.1038/nmeth.4605

55. Grimm JB, Klein T, Kopek BG, Shtengel G, Hess HF, Sauer M, et al. Synthesis of a far-red photoactivatable silicon-containing rhodamine for super-resolution microscopy. Angewandte Chemie Int Ed Engl. 2016;55: 1723–1727. doi:10.1002/anie.201509649

56. Grimm JB, Klein T, Kopek BG, Shtengel G, Hess HF, Sauer M, et al. Synthesis of a Far-Red Photoactivatable Silicon-Containing Rhodamine for Super-Resolution Microscopy. Angewandte Chemie. 2015;128: 1755–1759. doi:10.1002/ange.201509649

57. Parthasarathy R. Rapid, accurate particle tracking by calculation of radial symmetry centers. Nat Methods. 2012;9: 724–726. doi:10.1038/nmeth.2071

58. Ma H, Xu J, Jin J, Huang Y, Liu Y. A Simple Marker-Assisted 3D Nanometer Drift Correction Method for Superresolution Microscopy. Biophysical Journal. 2017;112: 2196–2208. doi:10.1016/j.bpj.2017.04.025

59. Tokunaga M, Imamoto N, Sakata-Sogawa K. Highly inclined thin illumination enables clear single-molecule imaging in cells. Nat Methods. 2008;5: 159–161. doi:10.1038/nmeth1171

60. Juette MF, Gould TJ, Lessard MD, Mlodzianoski MJ, Nagpure BS, Bennett BT, et al. Three-dimensional sub–100 nm resolution fluorescence microscopy of thick samples. Nat Methods. 2008;5: 527–529. doi:10.1038/nmeth.1211

61. Pavani SRP, Thompson MA, Biteen JS, Lord SJ, Liu N, Twieg RJ, et al. Three-dimensional, single-molecule fluorescence imaging beyond the diffraction limit by using a double-helix point spread function. Proc National Acad Sci. 2009;106: 2995–2999. doi:10.1073/pnas.0900245106

62. Harris R. Chemistry Nobel Given To Scientists For Work On Optical Microscope. NPR; 2014. Available: https://www.npr.org/2014/10/08/354639749/chemistry-nobel-given-to-scientists-for-work-on-optical-microscope

63. Coffman EN. Essays on PhD Output at U.S Undergraduate Institutions. 2012.

